# Dispersal-dependent juvenile survival in a sexually dimorphic long-lived bird, the Greater Flamingo *Phoenicopterus roseus*

**DOI:** 10.1101/368068

**Authors:** Guillaume Souchay, Christophe Barbraud, Christophe Germain, Arnaud Béchet

**Author notes:** ONCFS – DRE, Unité Petite Faune de Plaine, 8 Bd Albert Einstein CS 10 42355, 44323 Nantes Cedex 3, France.

## Abstract

The viability and dynamics of spatially structured populations depend critically upon dispersal behaviour. Yet, in long lived species with delayed maturity, the fitness consequences of post-fledging dispersal, dispersal from the birthplace after independence and before first breeding attempt, are poorly understood although it is a critical determinant of natal dispersal.

We aimed at estimating sex-specific variations of juvenile survival in a long-lived bird species with sexual size dimorphism, the greater flamingo, as a function of post fledging dispersal destination. Using capture-recapture models, we estimated the survivorship of flamingos ringed in the Camargue (south of France) and wintering in the Mediterranean.

Dispersal probability from France was > 0.66 with important annual variations in preferred dispersal destinations. First-year survival increased along the winter temperature gradient with estimates below 0.50 ± 0.07 in France and above 0.60 ± 0.07 in African wetlands. The survival of flamingos wintering in France dropped by 30–50 % depending on sex between fall and spring of their first year.

In African sites and in Italy, there was no detectable difference of survival between sexes suggesting favorable wintering conditions. Body condition at fledging did not explain variations in first-fall survival within genders. Males wintering in France had a better survival than females.

These results show that sex and post-fledging dispersal destination affect juvenile survival, support the *energetic* hypothesis predicting an advantage of large body size to cope with cold temperatures and the *competition* hypothesis, and suggest the existence of sex-specific post fledging dispersal tactics.

## Introduction

Dispersal is a key process in population dynamics and genetics (Clobert et al. 2001). Whereas most studies on dispersal have investigated the causes and consequences of natal and breeding dispersal (Greenwood 1980, Clobert et al. 2009), the fitness pay-offs of post-fledging dispersal remain poorly evaluated (but see Naef-Daenzer et al. 2001, Wiens et al. 2006). Post-fledging dispersal is the dispersal of the juveniles of both sexes from parental territories or from the birthplace during the post-fledging period, soon after they achieve independence (Morton et al. 1991). Post-fledging dispersal is likely to be a critical determinant of natal dispersal as it permits prospecting for potential future breeding sites (Bell 1991, Clobert et al. 2001).

Both exogenous and endogenous factors have been suggested to drive dispersal, including conspecific interactions (Dobson 1982, Greenwood and Harvey 1982, Waser 1985, Wiggett and Boag 1993, Balkız et al. 2010), declining food availability (Kenward et al. 1993), body condition (Belthoff and Dufty 1998, Clobert et al. 2001, Barbraud et al. 2003), experience (Balkız et al. 2010) or genetic determinants (Fleischer et al. 1984, Clobert et al. 2001). In these views, dispersal is a state-dependent behavioral response supposedly immediately beneficial in terms of improved body condition or increased survival.

Estimating sex-specific variations of juvenile survival as a function of both body condition and post-fledging dispersal destinations should help identifying fitness benefits of dispersal. This type of study is however hampered by several difficulties. In long-lived species, recruitment can be delayed for several years (Newton 1989, Bradley and Wooler 1991, Clobert and Lebreton 1991, Croxall and Rothery 1991, Frederiksen and Bregnballe 2000, Bogdanova et al. 2007, Harris et al. 2007) and for several species, young individuals do not attempt to breed at their first visit to the breeding area (Bradley and Wooler 1991), which usually results in a lower resighting probability. Hence in these species, dispersing juveniles are only resighted on breeding sites several years after they were marked as juveniles, making juvenile survival difficult to estimate properly: individuals could have died as soon as they left their birth place or just before their first return to the breeding area.

Here we estimate sex-specific first-year survival of a long-lived sexually dimorphic species, the greater flamingo *Phoenicopterus roseus* while taking into account possible modulation by body condition and dispersal destination. The greater flamingo can live up to 40 year-old in the wild and > 60 year-old in captivity (Johnson and Cézilly 2007). Natal and breeding dispersal are important in this species so that the breeding colonies of the Mediterranean function like a metapopulation *sensu lato* (Barbraud et al. 2003, Balkız et al. 2007b, Balkız et al. 2010). Post fledging dispersal of chicks born in the Camargue, Southern France, is above 60% (Barbraud et al. 2003) with fledglings observed all around the Mediterranean basin and in West Africa in their first winter. While previous studies had shown that post-fledging dispersal destination was affected by predominant wind directions at the time of departure (Green et al. 1989), the respective pay-offs of arrival destinations are unknown. These post-fledging wintering areas (Fig. 1) are characterised by a gradient of increasing winter temperatures from the south of France (average of the minimal values of the coldest winter months = 0–2°, Emberger et al., 1963) to Spain (4–6°), Italy (3–7°), North-Africa and Turkey (4–9°).

**Figure 1.**
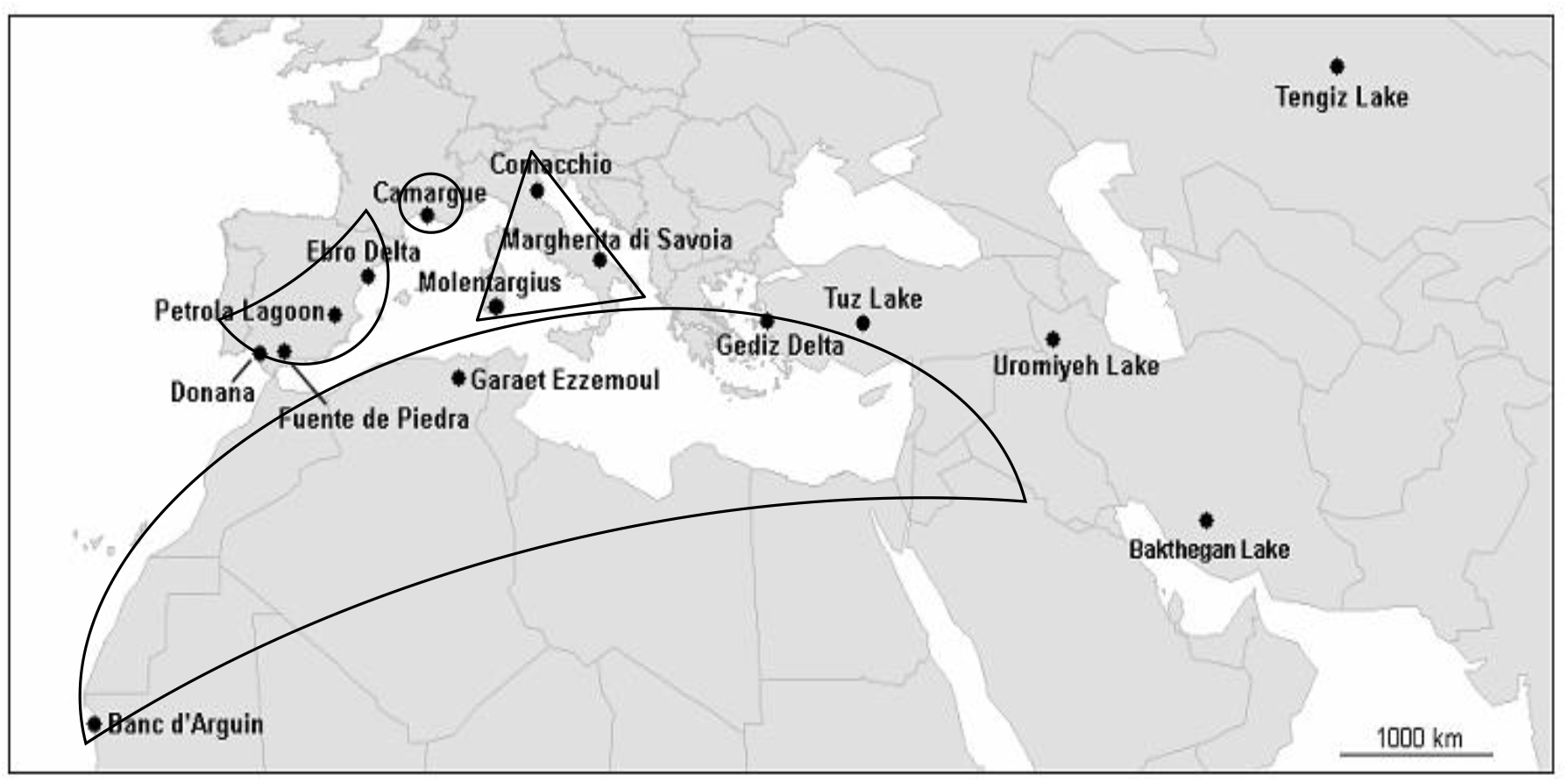
Colonies of greater flamingos (*Phoenicopterus roseus*) where breeding occurred at least once between 2000 and 2005 in the Mediterranean, West Africa and the Middle East. Regions used in the analysis are delimited by a dotted line

Predictions about the expected benefits of dispersal in term of survival in species presenting a sexual size dimorphism can be derived from hypotheses used to explain both differential dispersal and migration among classes of individuals (e.g. body size, sex or age; Greenwood 1980, Cristol et al. 1999):

### Prediction 1

According to the *body size* (or *energetic*) hypothesis, differential movements might result if population classes differ in body size to such an extent that it affects ability to survive poor environmental conditions. In one hand, larger individuals would be more likely than smaller individuals to survive the cold temperatures because of a more favorable surface-area-to-volume ratio (Ketterson and Nolan 1976). In the other hand, food shortage resulting from competition and/or poor environmental conditions may affect more seriously the larger individuals due to their higher energetic requirements for maintenance (Clutton-Brock et al. 1985, Demment and Vansoest 1985, Promislow 1992, Toïgo and Gaillard 2003, Kitaysky et al. 2006, Harris et al. 2007). Adult male greater flamingos being 20% taller and heavier than females (Johnson and Cézilly 2007), we thus predict female juvenile survival to differ from male juvenile survival in the south of France, the colder wintering area. If body size is an advantageous character in cold climate, we consequently expect males to perform better in this region. Alternatively, if body size is a handicap, males should be disadvantaged.

### Prediction 2

According to the *dominance* hypothesis, dominants occupy either sites of better quality or closest to the breeding sites, subordinates being displaced farther away from the breeding grounds or from high quality areas (Gauthreaux Jr 1978). This hypothesis assumes inequalities in resource holding potential between individuals of one class (e.g. large individuals) that can limit access to resources to another class (e.g. smaller individuals:Hunter 1983, Rowland 1989, Langston et al. 1990). It also makes the assumption that movements are costly (Cristol et al. 1999). According to the *competition* hypothesis, our prediction would be that male juvenile survival is higher than female juvenile survival in sites of better quality. This would result from males dominating females on foraging patches of good quality displacing them to low quality wintering areas. This would for instance be the case in the south of France which hold one of the largest colonies of the species in the Mediterranean and present increasing numbers of wintering flamingos (Johnson and Cézilly 2007; Béchet et al. unpublished data). In this case, female survival would increase with increasing distance to the colony by decreasing competition.

### Prediction 3

Survival variations may be modulated by individual variations in body condition. Indeed, survival of migratory birds was proven to be condition-dependent with individuals in better condition benefitting from increased survival (Sandberg and Moore 1996, Bety et al. 2003, Schekkerman et al. 2003, Morrison et al. 2007). Hence, based on previous findings in the greater flamingos (Barbraud et al. 2003) and other species (Magrath 1991, Clobert et al. 2001, Naef-Daenzer et al. 2001, Menu et al. 2005, Braasch et al. 2009), we expect a positive effect of body condition on juvenile survival within genders irrespective of the dispersal destination.

## Methods

### Species and study area

Greater flamingos are seasonally monogamous birds (Cézilly and Johnson 1995) inhabiting temporary brackish or salt marshes (Johnson and Cézilly 2007, Bechet et al. 2009). Since 1969, flamingos have been breeding every year (except 2007) at the Fangassier lagoon (43°25’N, 4°37’E) in the Camargue, south of France, with an average of 10 000 breeding pairs (Bechet et al. 2011).

From 1995 to 2005, 3 519 chicks have been marked with PVC rings and sexed genetically (Bertault et al. 1999, see Balkız et al. 2007a and Supplementary Material Table S1) just before fledging. Body mass and tarsus length were recorded at ringing. Tarsus length was measured (± 1 mm) as the distance from the distal end of tarsometarsus to the outer edge of the midtarsal joint (Table S2). Body mass was measured to the nearest 50 g using a 0–5000 g Pesola spring balance (Barbraud et al. 2003).

Ring codes can be read using a telescope and from a distance up to 300 meters (Johnson and Cézilly 2007). We divided the resighting area (Fig. 1) into four regions considered as the most important wintering sites for the species in the Mediterranean: southern France (Fr), Spain (from the Ebro delta to Andalusia; Sp), Italy (continental Italy and Sardinia; It), African and Turkish sites (from Mauritania to Turkey; Ot). This last region encompasses a large diversity of wetlands where observation effort is low.

### Data analysis

We focused on the resightings of these ringed flamingos from winter 1995 to summer 2007. In order to estimate separately fall and spring survival, we divided each year into two resighting sessions: *Summer*, which includes resightings made from April to July, and *Winter*, which includes resightings made from November to January.

### Modelling resighting and movement probabilities

We used Jolly-MoVement (JMV) multisite models (Lebreton et al. 2009) to estimate movement and survival probabilities conditional on site-specific resighting probabilities. Because we aimed at estimating sex-specific juvenile survival we constrained resightings and movement probabilities based on previous findings. In particular, movements were constrained according to Barbraud et al. (2003) who found dispersal to be time-dependent for the first age class (birds < 1 year old) and time-independent for the second age-class (birds aged ≥ 1 year old). Because we were interested in estimating seasonal survival probability, we considered individuals < 6 month-old as a first age-class, and individuals > 6 month-old (from 6 month-old to 12 month-old) as the second age-class.

Previous demographic studies on greater flamingos from the Camargue (Lebreton et al. 1992, Cézilly et al. 1996, Tavecchia et al. 2001, Barbraud et al. 2003) found that resighting probabilities varied according to year, age and site. Therefore, in our initial model, resighting probabilities were site-, time- and age-specific, with the same age-classes as those used for movement probabilities.

### Goodness-of-fit test and model selection

The ability of the initial model to describe the data was assessed by goodness-of-fit tests run for each cohort and for each sex using U-CARE (Choquet et al. 2009a). We then followed a step down approach, starting with the initial model and fitting models with constrained parameterizations for resighting and survival probabilities. Model selection relied on the Akaike’s Information Criterion (AIC, Burnham and Anderson 2002). Model selection and parameter estimation were performed using M-SURGE (Choquet et al. 2004), except for the effect of body condition which was modelled with E-SURGE(Choquet et al. 2009b), as this software enable the use of individual covariates. We evaluated the possible improvement of the best model by modelling first-year survival as a function of body condition. All values are given ± SE.

### Modelling survival probabilities

In our initial model adult survival was age-, time- and site-specific. Since we were interested in variations of survival probabilities early in life, we defined two age-classes for the first year of life: first-fall (i.e. from ringing in July to January), and first-spring (i.e. from January to July the following year). After the first year of life we allowed survival probabilities to vary with age on an annual basis until age 3, and independently of age afterward since previous studies suggested that survival probabilities varied little according to age after age 3. We also allowed survival to be different before and after age 7, the peak of recruitment, for some models (Johnson et al. 1991, Lebreton et al. 1992, Cézilly et al. 1996, Tavecchia et al. 2001, Johnson and Cézilly 2007).

Using the notation recommended by Lebreton and Pradel (2002) our initial model was therefore: 
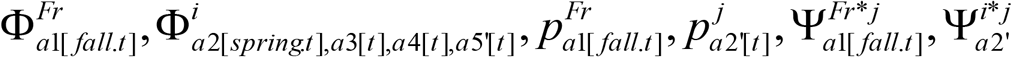
where Φ, *p* and Ψ are survival, resighting and movement probabilities respectively, *i* and *j* the departure and current site (Fr, Sp, It, or Ot), *t* the resighting occasions – time effect is the interaction between years and seasons, *spring* and *fall* respectively indicate first post-winter and first post-summer periods, and *a#* indicates age-classes (see Table S3 for the details of shortcuts used).

Movement probabilities for the first age-class differ from subsequent ones as all birds were ringed in France so that France is the only possible site of departure for this age-class. For birds belonging to older age-classes movement probabilities were site-specific but were set constant with time to limit the number of parameters in the model relative to the available dataset. As all birds were ringed in the Camargue, survival probability of the first age-class was only estimated for the Camargue. For this first age-class, survival and resighting probabilities for Spain, Italy and Others were fixed to 1 as no birds of this age were available at these sites. For the subsequent age-classes, resighting probabilities were only site- and time-dependent, and survival probabilities were age- and time-dependent until age 3, and only time-dependent for the older age-classes.

The power and memory requirements of the software M-SURGE used to run the models (Choquet et al. 2004) were exceeded to run a model including a sex effect on survival probabilities (4 sites, 4 age classes, 2 sexes, seasonal variation corresponding here to the number of occasions, 3 519 individuals and *c*. 3 600 resightings). We thus divided the data set into two, according to sex, and compared *a posteriori* sex-specific survival estimates using Z-tests (Zar 1984). We used Gould and Nichols’ method (1998) to estimate process variance when temporal variation was detected. We could then compare average parameter estimates (survival and movement probabilities) using Z-tests. When multiple comparisons were performed *p*-values were Bonferroni corrected (*pBonf* = *p*-values * number of comparisons).

### Body condition effect on survival

A body condition index (BCI) was estimated for each individual as the residual from a reduced major axis regression of the log of body mass on the log of tarsus length for each year (Green 2001, Barbraud et al. 2003). Residuals were calculated with the Smatr package in program R (Warton 2007, R Development Core Team 2010) for 1 592 males and 1 914 females. Residuals were standardized and used as an individual covariate when testing for an effect of body condition on juvenile survival.

## Results

### Goodness-of-fit tests

There was no lack of fit for the JMV model when data were pooled by sex and cohort (Table S4). The overall goodness of fit test suggested an underdispersion of the data as *ĉ* was < 1 for both sexes. Underdispersion likely resulted from a low number of resightings per bird (0.96 ±0.04 observations per female and 1.08 ± 0.04 observations per male). As suggested by Cooch and White (2007) we thus kept a value of 1 for the ĉ.

### Model selection

We started by modelling the male dataset. Starting with our initial model (model P2; Table S5), we examined whether resighting probabilities were age-, year-, season-, and site-dependent (models P1 to P6; Table S5). The full time-dependent model was preferred indicating that resighting probabilities varied between seasons, years and sites irrespective of age.

Regarding survival, an age-dependent model (S2) was preferred to a year-dependent model (S4) or to a year- and age-dependent model (S7), suggesting that survival did not vary among years (Table 1). A model with survival differing between juveniles, immatures and adults (S1), was preferred to a model where survival differed between juveniles, immatures, young and old adults (S3) suggesting no detectable age variations in survival after age 3. Hence, this best model (S1) distinguished 4 age-classes: from fledging to 1^st^ fall, from 1^st^ fall to 1^st^ spring, from 1 to 2 year-old, and ≥ 3 year-old.

**Table 1.**
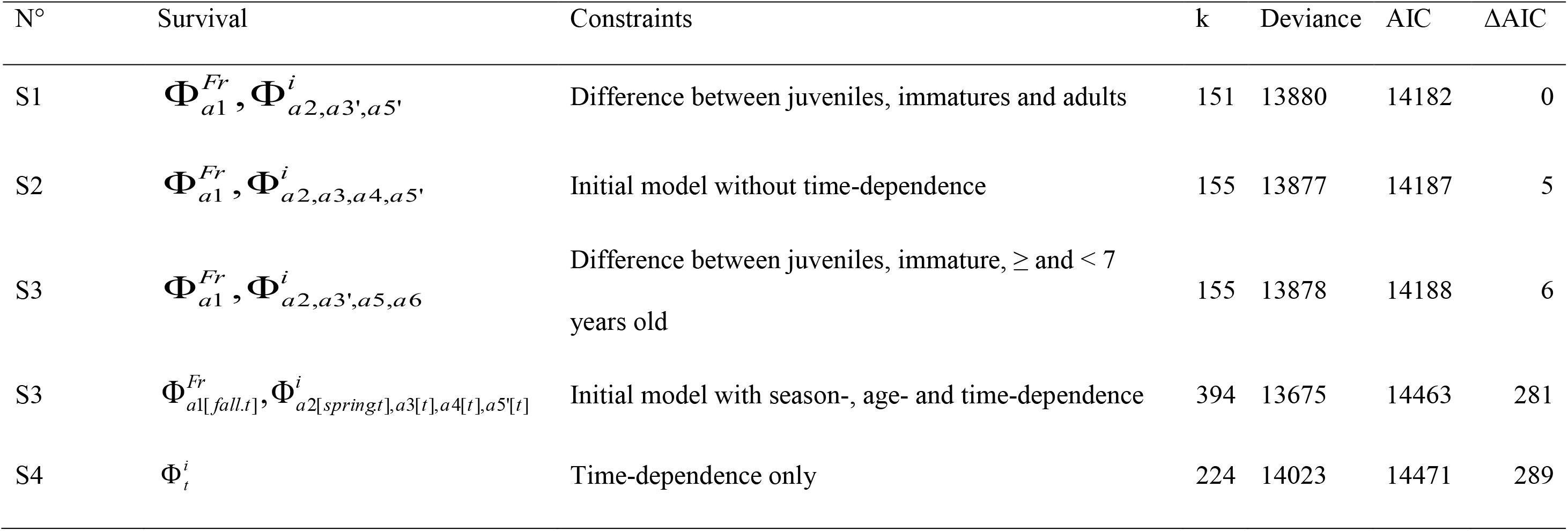
Modelling male survival probability Φ of Greater flamingos born in the Camargue, France between 1995 and 2005. Resighting probabilities were modelled as 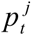 and movement probabilities as 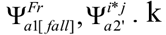 (see Methods for model notation). *k* is the number of estimated parameters. Models are sorted by ΔAIC values.

As for males, female resighting probabilities were site-, season- and time-dependent (Table S6). The same model structure as for males was selected for survival (Table 2).

**Table 2.** Modelling female survival probability Φ of greater flamingos born in the Camargue, France between 1995 and 2005. Resighting probabilities were modelled as 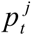 and movement probabilities were modelled as 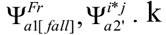 (see Methods for model notation). k is the number of estimated parameters. Models are sorted by ΔAIC values.

| N° | Survival | Constraints | k | Deviance | AIC | $\Delta AIC$ |
| --- | --- | --- | --- | --- | --- | --- |
| S1' | $\Phi_{a1}^{Fr}, \Phi_{a2,a3',a5'}^i$ | Difference between juveniles, immatures and adults | 151 | 14960 | 15262 | 0 |
| S2' | $\Phi_{a1}^{Fr}, \Phi_{a2,a3,a4,a5'}^i$ | Initial model without time-dependence | 155 | 14958 | 15268 | 6 |
| S3' | $\Phi_{a1}^{Fr}, \Phi_{a2,a3',a5,a6}^i$ | Difference between juveniles, immature, ≥ and $< 7$<br>years old | 155 | 14960 | 15270 | 8 |
| S7' | $\Phi_t^i$ | Time-dependence only | 225 | 15063 | 15513 | 251 |
| S8' | $\Phi_{a1[fall,t]}^{Fr}, \Phi_{a2[springt],a3[t],a4[t],a5[t]}^i$ | Initial model with season-, age- and time-dependence | 393 | 14759 | 15545 | 283 |

### Parameters estimates

#### Movement probabilities

Post-fledging dispersal was high with less than 30 % of fledglings remaining in the south of France for their first winter (Ψ^Fr-Fr^ < 0.30 for both sexes). African and Turkish sites were the preferred post-fledging destinations (range 0.11 ± 0.05 to 0.77 ± 0.22) followed by Spain and Italy (Table 3). During their first-year, young male flamingos had the highest probability to stay in France (0.153 ± 0.003 against 0.126 ± 0.011for females, z-test: *z*= 2.4510, *p* = 0.014), and the highest probabilities to go to Spain (0.308 ± 0.018 against 0.208 ± 0.002, *z*= 5.6736, *p* < 0.001) and to Italy (0.149 ± 0.006 against 0.120 ± 0.001, *z*= 4.9919, *p* < 0.001) while females had the highest post-fledging movements to African and Turkish sites (0.546 ± 0.020 against 0.389 ± 0.038, *z*= 3.6642, *p* < 0.001). As expected, subsequent movements were low confirming a high philopatry to the wintering site in this species (Ψ^i-i^ > 0.86 for both sexes).

#### Survival probabilities

Because survival estimates were high from the second fall onward (> 0.90 for all sites and both sexes, Table 3 and Table S7 for annual survival), we focused only on the first-year estimates (Fig. 2).

**Figure 2.**
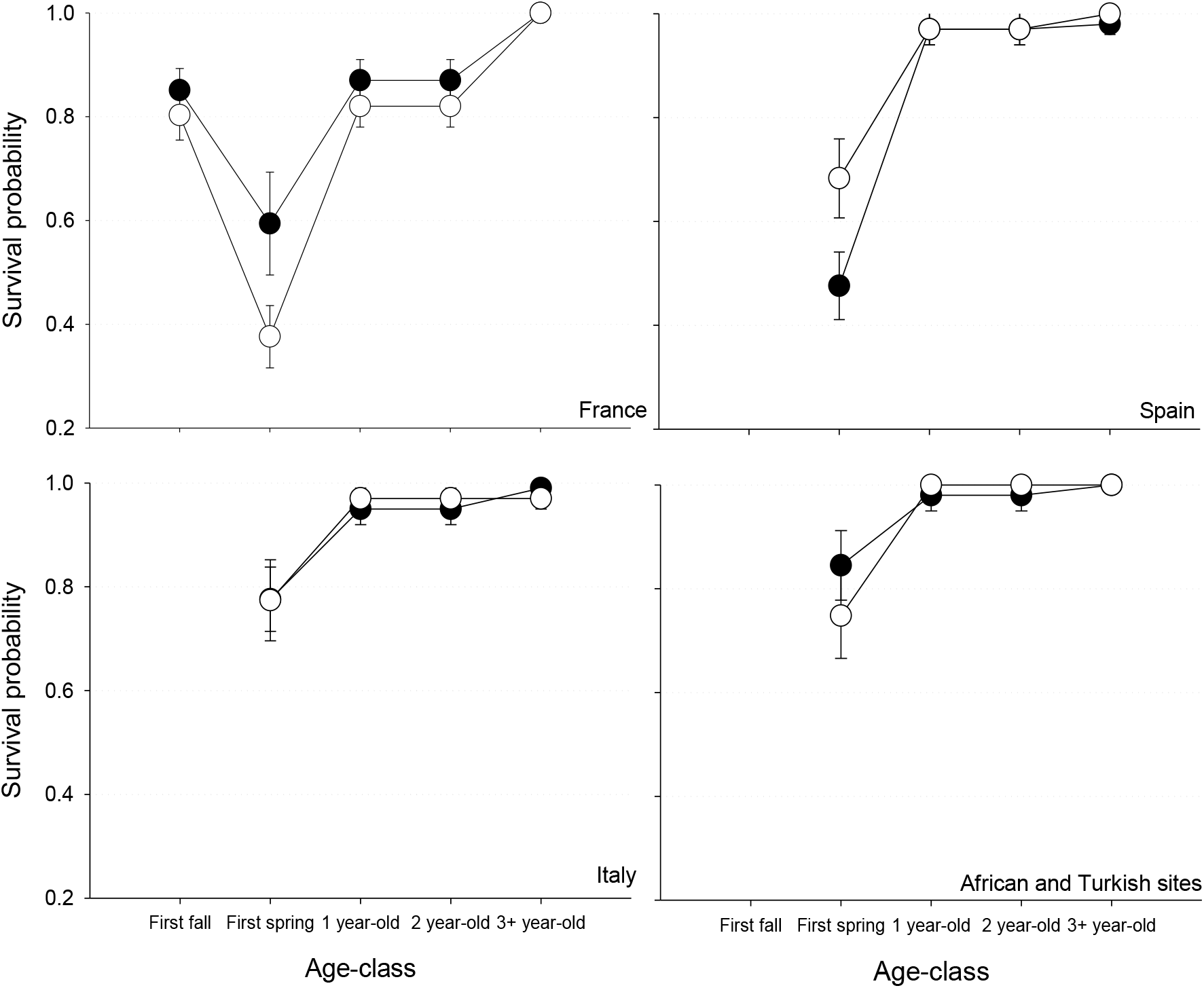
Survival probabilities (±SE) for males (filled circle) and females (open circle) greater flamingos born from 1995 to 2005 and resighted from 1995 to 2007 in France, Spain, Italy, Africa and Turkey. Probabilities were estimated from models S1 and S1’ for males and females, respectively

**Table 3.**
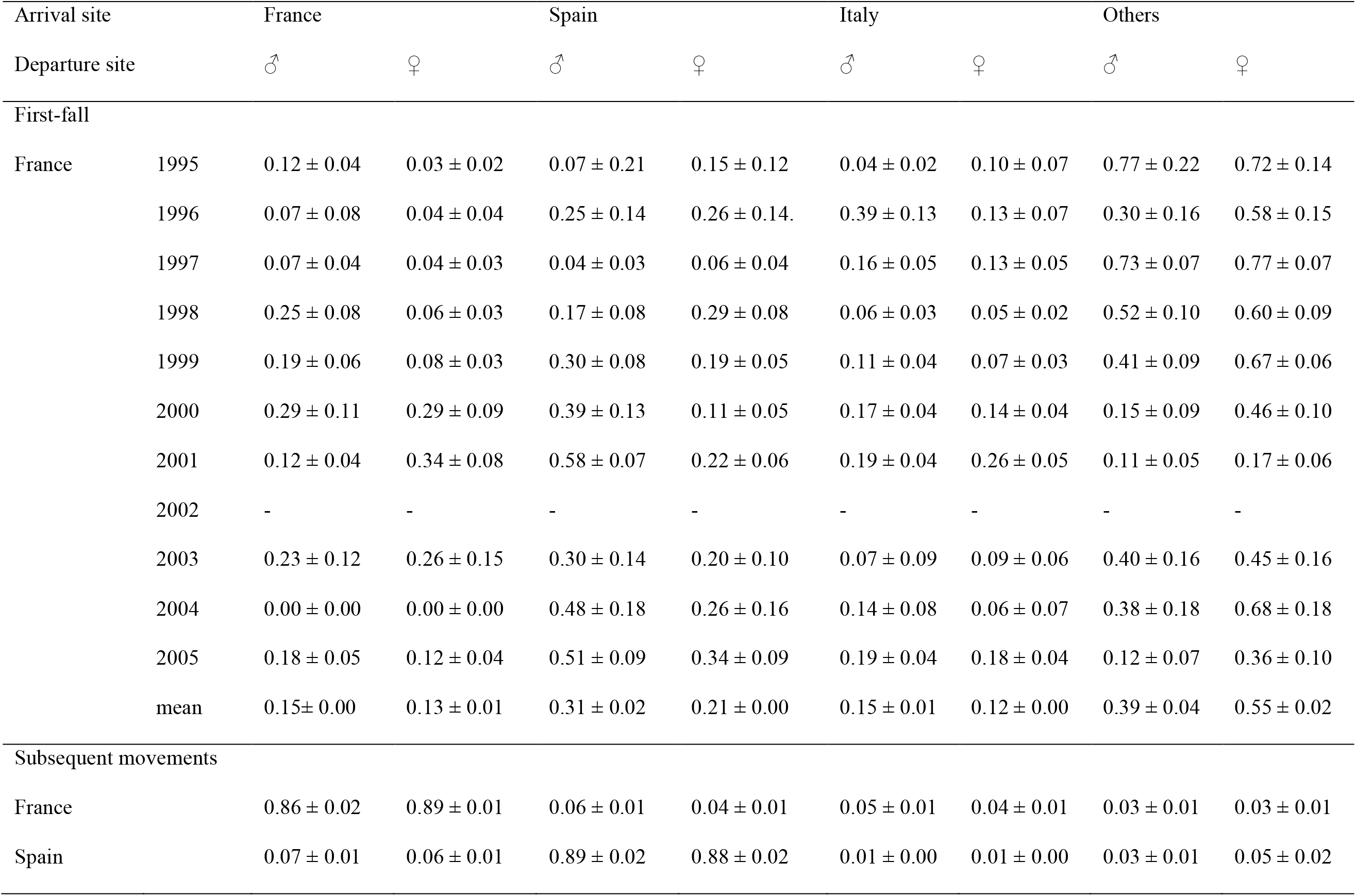
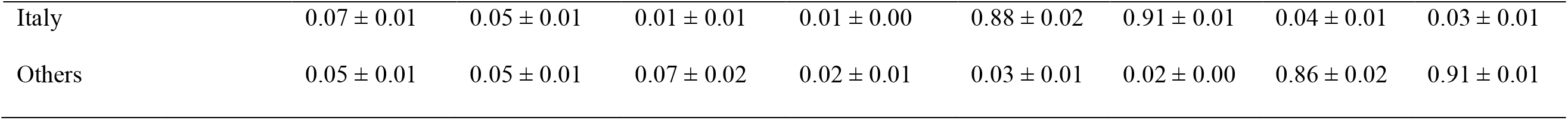
Movement probabilities (± SE) for Greater flamingos born in the Camargue between 1995 and 2005 by sites from models S1 and S’1.

| Arrival site |  | France |  | Spain |  | Italy |  | Others |  |
| --- | --- | --- | --- | --- | --- | --- | --- | --- | --- |
| Departure site |  | ♂ | ♀ | ♂ | ♀ | ♂ | ♀ | ♂ | ♀ |
| First-fall |  |  |  |  |  |  |  |  |  |
| France | 1995 | 0.12 ± 0.04 | 0.03 ± 0.02 | 0.07 ± 0.21 | 0.15 ± 0.12 | 0.04 ± 0.02 | 0.10 ± 0.07 | 0.77 ± 0.22 | 0.72 ± 0.14 |
| | 1996 | 0.07 ± 0.08 | 0.04 ± 0.04 | 0.25 ± 0.14 | 0.26 ± 0.14 | 0.39 ± 0.13 | 0.13 ± 0.07 | 0.30 ± 0.16 | 0.58 ± 0.15 |
| | 1997 | 0.07 ± 0.04 | 0.04 ± 0.03 | 0.04 ± 0.03 | 0.06 ± 0.04 | 0.16 ± 0.05 | 0.13 ± 0.05 | 0.73 ± 0.07 | 0.77 ± 0.07 |
| | 1998 | 0.25 ± 0.08 | 0.06 ± 0.03 | 0.17 ± 0.08 | 0.29 ± 0.08 | 0.06 ± 0.03 | 0.05 ± 0.02 | 0.52 ± 0.10 | 0.60 ± 0.09 |
| | 1999 | 0.19 ± 0.06 | 0.08 ± 0.03 | 0.30 ± 0.08 | 0.19 ± 0.05 | 0.11 ± 0.04 | 0.07 ± 0.03 | 0.41 ± 0.09 | 0.67 ± 0.06 |
| | 2000 | 0.29 ± 0.11 | 0.29 ± 0.09 | 0.39 ± 0.13 | 0.11 ± 0.05 | 0.17 ± 0.04 | 0.14 ± 0.04 | 0.15 ± 0.09 | 0.46 ± 0.10 |
| | 2001 | 0.12 ± 0.04 | 0.34 ± 0.08 | 0.58 ± 0.07 | 0.22 ± 0.06 | 0.19 ± 0.04 | 0.26 ± 0.05 | 0.11 ± 0.05 | 0.17 ± 0.06 |
|  | 2002 | - | - | - | - | - | - | - | - |
| | 2003 | 0.23 ± 0.12 | 0.26 ± 0.15 | 0.30 ± 0.14 | 0.20 ± 0.10 | 0.07 ± 0.09 | 0.09 ± 0.06 | 0.40 ± 0.16 | 0.45 ± 0.16 |
| | 2004 | 0.00 ± 0.00 | 0.00 ± 0.00 | 0.48 ± 0.18 | 0.26 ± 0.16 | 0.14 ± 0.08 | 0.06 ± 0.07 | 0.38 ± 0.18 | 0.68 ± 0.18 |
| | 2005 | 0.18 ± 0.05 | 0.12 ± 0.04 | 0.51 ± 0.09 | 0.34 ± 0.09 | 0.19 ± 0.04 | 0.18 ± 0.04 | 0.12 ± 0.07 | 0.36 ± 0.10 |
| | mean | 0.15 ± 0.00 | 0.13 ± 0.01 | 0.31 ± 0.02 | 0.21 ± 0.00 | 0.15 ± 0.01 | 0.12 ± 0.00 | 0.39 ± 0.04 | 0.55 ± 0.02 |
| Subsequent movements |  |  |  |  |  |  |  |  |  |
| France | | 0.86 ± 0.02 | 0.89 ± 0.01 | 0.06 ± 0.01 | 0.04 ± 0.01 | 0.05 ± 0.01 | 0.04 ± 0.01 | 0.03 ± 0.01 | 0.03 ± 0.01 |
| Spain | | 0.07 ± 0.01 | 0.06 ± 0.01 | 0.89 ± 0.02 | 0.88 ± 0.02 | 0.01 ± 0.00 | 0.01 ± 0.00 | 0.03 ± 0.01 | 0.05 ± 0.02 |

|  |  |  |  |  |  |  |  |  |
| --- | --- | --- | --- | --- | --- | --- | --- | --- |
| Italy | $0.07 \pm 0.01$ | $0.05 \pm 0.01$ | $0.01 \pm 0.01$ | $0.01 \pm 0.00$ | $0.88 \pm 0.02$ | $0.91 \pm 0.01$ | $0.04 \pm 0.01$ | $0.03 \pm 0.01$ |
| Others | $0.05 \pm 0.01$ | $0.05 \pm 0.01$ | $0.07 \pm 0.02$ | $0.02 \pm 0.01$ | $0.03 \pm 0.01$ | $0.02 \pm 0.00$ | $0.86 \pm 0.02$ | $0.91 \pm 0.01$ |

First-fall survival was high and did not differ between sexes (0.85 ± 0.04 and 0.80 ± 0.05, for males and females respectively, *z* = 0.75, *p* = 0.45).

First-spring survival of male flamingos staging in France (0.59 ± 0.10) did not differ significantly from those which had dispersed (Table 4; all z-test paired comparisons: *p*> 0.05). However, among dispersers, males had a lower survival rate in Spain (0.48 ± 0.07) than in Italy (0.78 ± 0.06) and in African and Turkish sites (0.84 ± 0.07; z*Sp/It* = 3.33 *pBonf* < 0.006; *zSp/Ot* = 3.93, *pBonf* < 0.001). Female first-spring survival (0.38 ± 0.06) was much lower in France than everywhere else (>0.68; *zFr/Sp* = 3.17, *pBonf* < 0.01, *zFr/It*= 4.03, *pBonf* < 0.001, *z* Fr/Ot = 3.67, *pBonf* = 0.0012). There was no site variation in first-spring survival probabilities between female dispersers.

**Table 4.**
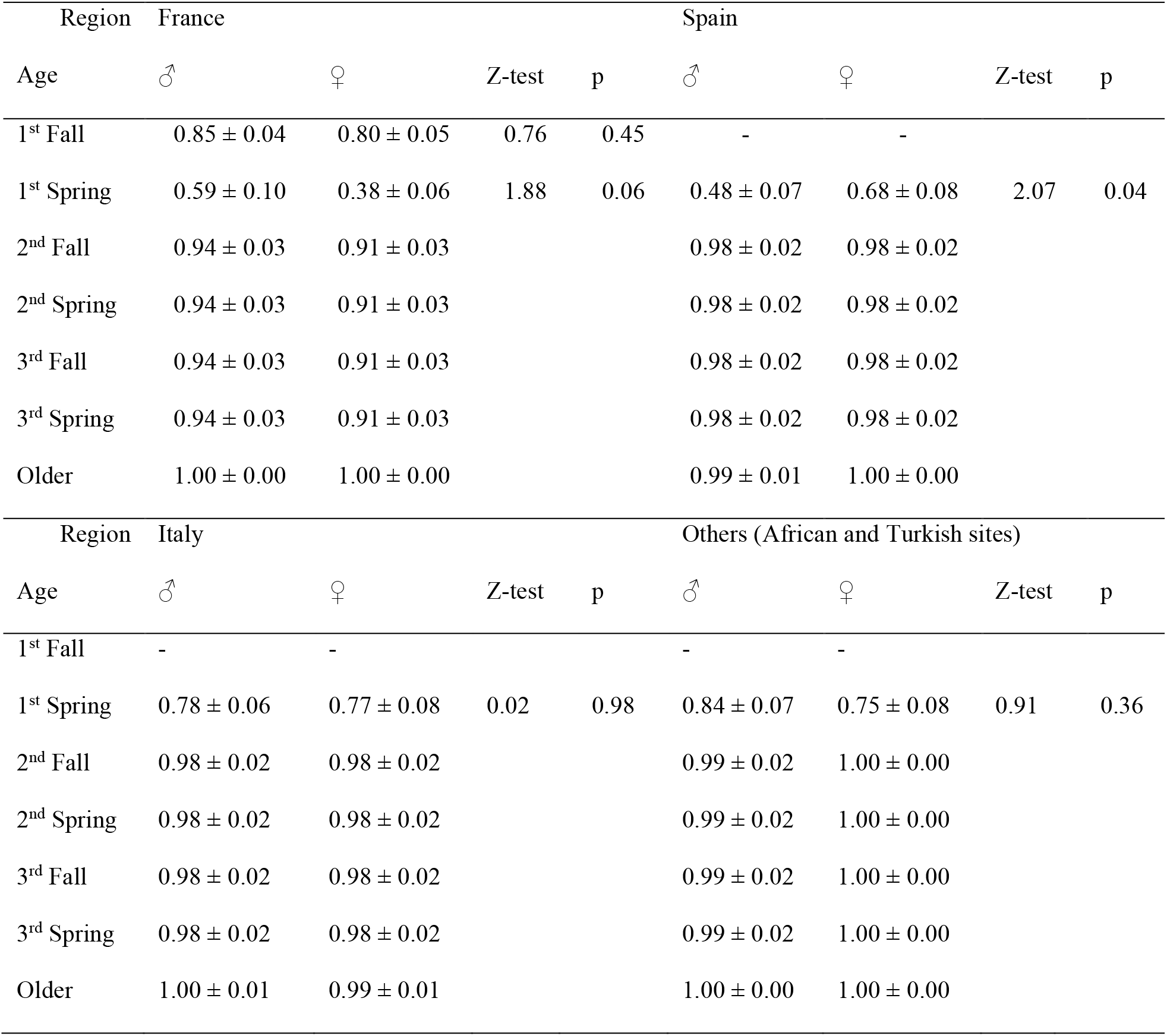
Survival probabilities (± SE) of Greater flamingos born in the Camargue between 1995 and 2005 by sex and by sites from models S’1 and S1. Survival differences between sexes and by sites are tested by Z-tests.

In France, there was a strong tendency for males to have a higher first-spring survival than females (0.59 ± 0.04 *vs*. 0.38 ± 0.06, *z* = 1.88, *p* = 0.06). For dispersers, female and males had similar survival probabilities except in Spain where females appeared to do better than males (0.48 ± 0.07 for males *versus* 0.68 ± 0.08 for females, *z* = 2.07, *p* = 0.04).

#### Body condition effect on survival

Sexual dimorphism was apparent before fledging as male chicks were on average 7.3% larger and 13% heavier than females, respectively (tarsus length: *t*= 13.26, df = 3509, *p* < 0.001; body mass: *t*= 12.71, df = 3505, *p* < 0.001; Table S2).

For males, a model forcing survival to depend on body condition was only marginally better supported than a model without covariate (ΔDeviance = 10 but ΔAIC <2, Table 5). Furthermore, slopes were not different from zero. For females, a model including the effect of body condition on survival probabilities did not improve the AIC (Table 6).

**Table 5.** Modelling the effect of body condition on male survival probability Φ with resighting parameter 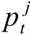 and movement parameter 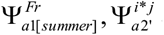.k is the number of estimated parameters. Models are sorted by ΔAIC values.

| Model | Constraints | k | Deviance | AIC | $\Delta AIC$ |
| --- | --- | --- | --- | --- | --- |
| $\Phi_{a1[BCI]}^{Fr}, \Phi_{a2[BCI],a3',a5'}^i$ | BCI effect on both first-fall and first-spring survival | 156 | 13,810 | 14,122 | 0 |
| $\Phi_{a1}^{Fr}, \Phi_{a2[BCI],a3',a5'}^i$ | BCI effect on first-spring survival | 156 | 13,811 | 14,123 | 1 |
| $\Phi_{a1}^{Fr}, \Phi_{a2,a3',a5'}^i$ | Initial model without BCI effect | 151 | 13,821 | 14,123 | 1 |
| $\Phi_{a1[BCI]}^{Fr}, \Phi_{a2,a3',a5'}^i$ | BCI effect on first-fall survival | 152 | 13,820 | 14,124 | 2 |
| $\Phi_{a1}^{Fr}, \Phi_{a2+BCI,a3',a5'}^i$ | Additive effect of BCI on first-spring survival | 152 | 13,821 | 14,125 | 3 |

**Table 6.** Modelling the effect of body condition on female survival probability Φ with resighting parameter 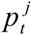 and movement parameter 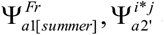.k is the number of estimated parameters. Models are sorted by ΔAIC values.

| Model | Constraints | k | Deviance | AIC | $\Delta AIC$ |
| --- | --- | --- | --- | --- | --- |
| $\Phi_{a1}^{Fr}, \Phi_{a2,a3',a5'}^i$ | Initial model without bci effect | 151 | 14,911 | 15,213 | 0 |
| $\Phi_{a1[BCI]}^{Fr}, \Phi_{a2,a3',a5'}^i$ | Bci effect on first fall survival | 152 | 14,911 | 15,215 | 2 |
| $\Phi_{a1}^{Fr}, \Phi_{a2[BCI],a3',a5'}^i$ | Bci effect on first spring survival | 155 | 14,908 | 15,218 | 5 |
| $\Phi_{a1[BCI]}^{Fr}, \Phi_{a2[BCI],a3',a5'}^i$ | Bci effect on both first fall and first<br>spring | 156 | 14,906 | 15218 | 5 |

## Discussion

Our results confirm previous analyses showing that > 66% of greater flamingo fledglings disperse from the south of France (Barbraud et al. 2003) with important annual variations in preferred dispersal destinations reached for their first winter (Green et al. 1989). We also confirmed that flamingos presented a low dispersal (i.e. high wintering site fidelity) during subsequent years (Green et al. 1989, Barbraud et al. 2003).

The survival of greater flamingos during their first year increases along the winter temperature gradient with estimates below 0.50 ± 0.07 in France and above 0.60 ± 0.07 in North Africa and Turkey (Appendix Table S6). In France, survival of individuals in their first year drops by 30 to 50% depending on sex between fall and spring (Table 4). Therefore most of the mortality of juvenile greater flamingos born in the Camargue seems to occur between January and July of the year following their birth. This relatively heavy mortality rate may be caused by increasing energetic demands of fledgling flamingos facing colder temperatures accentuated by relatively strong winds during this period of the year in the south of France (Camargue). During their first-year, young birds progressively shift from parental care to autonomy and face the hazard of new surroundings, while gaining experience in obtaining suitable food resources (Anders et al., 1997, Kershner et al., 2004, Anders et al., 1998, Nilsson and Smith, 1985, Verhulst, 1992). Hence, in fall juveniles may benefit from parental care and appropriate body condition at fledging, while decreased survival in spring may reflect the cost of gaining foraging efficiency and perhaps decrease in food resources.

First-spring survival was the highest for both sexes in African and Turkish sites and in Italy. While African and Turkish sites constitute the preferred dispersal destination (up to 77% of the post-fledging dispersers), Italy does not attract more birds than Spain or France (Table 3). High first-spring survival in both African and Turkish sites and Italy (similar to first-fall survival in France), and similar survival between sexes suggest favorable wintering conditions at these sites. Relatively mild temperatures and availability of large wetlands in North and West Africa certainly offer appropriate conditions for flamingos in winter (Samraoui and Samraoui, 2008), while a small breeding population in Italy may provide a wintering area with low competition.

For birds remaining in France or dispersing to Spain (from 0 to 70% of the fledglings depending on years; Table 4), we found contrasted survival among sexes (Figure 2 and Appendix Table S6). The higher survival of males staying in France compared to females supports both the *energetic* hypothesis, predicting an advantage of a larger body size to cope with cold temperatures, and the *competition* hypothesis. According to the *competition* hypothesis, food limitation will advantage the more competitive larger individuals (Anderson et al., 1993, Rowland, 1989, Langston et al., 1990, Hunter, 1983, Arroyo, 2002) eventually causing higher risk of mortality for the smaller ones (Arroyo, 2002, Edwards and Collopy, 1983). This was found in the Montagu’s harrier (*Circus pygargus*) where male chicks weigh about 85% of females weight and showed a higher probability of starvation as a result of sibling competition (Arroyo, 2002). In blue tits (*Parus caeruleus*), a dimorphic species where males are bigger than females, Råberg et al. (2005) found that in poor conditions, female chicks suffer more heavily from starvation than male chicks. This was explained by the fact that female chicks, being smaller, were disadvantaged when competing for food delivered by parents (Råberg et al., 2005). Therefore, juveniles, and especially young females, may be more sensitive to competition and constrained to feed at poor quality sites in winter, when food is scarce and when the energetic needs increased with decreasing temperatures. This is especially likely for wintering sites situated at the northern edge of the greater flamingo range such as the south of France. In turn, dispersal to more temperate winter areas would be an advantageous tactic for juvenile females as suggested by the better juvenile survival observed elsewhere, and by the higher dispersal probability of females than males during their first year to African and Turkish sites.

An alternative explanation of the better survival of dispersing birds could be that dispersers are of better quality than sedentary ones. Altwegg et al. (, 2000) found that there was a tendency for larger House sparrows (*Passer domesticus*) to disperse farther than small ones. This would be consistent with the finding of Barbraud et al. (2003) showing that juvenile dispersal rates were higher with increased body condition, a possible proxy for individual quality. For subsequent age-classes and both sexes, survival remains the lowest in France, reinforcing the idea of a direct benefit of dispersal in term of survival. However, we only found a weak tendency for a positive effect of body condition on male juvenile survival, and no effect on female juvenile survival.

Previous studies had shown that flamingos wintering in France could be heavily affected by cold spells (Cézilly et al. 1996, Pradel et al., 1997, Tavecchia et al. 2001). Our results suggest that even in years without cold spell non-dispersing juveniles of both sexes pay the cost of a lower survival in the south of France. The maintenance of this tactic into the population may therefore result from other advantages of staying in France balancing the lower survival. Residents may have a better access to breeding sites as suggested by the time-allocation hypothesis (Greenberg, 1980). Despite attempts to specifically test this hypothesis on mallards (*Anas platyrhynchos* Hestbeck et al., 1992) or on sparrows (Sandercock and Jaramillo, 2002), it remains invalidated. Since in the greater flamingo there is only one egg laid and one reproduction attempt per year, we may hypothesize that an advantage could be a lower age of first-reproduction and/or a higher reproductive success. This remains to be tested.

Overall, considering first-year survival variations as a proxy for the immediate fitness payoffs of various post-fledging dispersal tactics, we can speculate on optimal sex-specific tactics. The best post-fledging dispersal tactic for females would be to move to temperate areas such as African and Turkish sites and Italy. Dispersing in Spain would be an intermediate option with intermediate payoffs, while remaining in France appears to be the most risky tactic. Observed post-fledging destinations of females respond rather well to this scoring as preferred destinations are the most advantageous in term of survival. Males should also preferentially disperse to African and Turkish sites, followed by Italy, France and Spain where winter appears harsh for this sex. However, as opposed to females, an important proportion of males seem to choose “looser” destinations so that the best and the worst options in term of survival are almost chosen equally (ψ^Ot^ = 0.389 ± 0.038 and ψ^Sp^ = 0.308 ± 0.018, respectively). Our results thus invite to explore other fitness advantages that could permit the maintenance of various post-fledging strategies in this species. In particular, as breeding experience was shown to be an important determinant of adult dispersal in the greater flamingo (Balkız et al. 2010), evaluating how post-fledging dispersal may favour earlier recruitment (and thus increased experience) at nearby foreign colonies could shed a new light for understanding the maintenance of such diverse dispersing strategies.

## Acknowledgment

This work is part of a long-term study of greater flamingos initiated by Dr Luc Hoffmann and supported by Tour du Valat and MAVA Foundation. We are grateful to the many people who participated in the fieldwork by sending their sightings of ringed flamingos. We thank Pierre Legagneux, Ana Sanz and Roger Pradel for helpful comments on analysis and earlier drafts of this manuscript.

## Supplementary materials

Table S1. Number of marked and sexed greater flamingos from 1995 to 2005 in the Camargue and resighted within the Mediterranean region from 1995 to 2007

Table S2. Mean ± SE (sample size) body mass and tarsus length for juvenile male and female greater flamingos ringed and sexed at the Camargue colony between 1995 and 2005.

Table S3. Shortcuts used in the definition of age-classes when modelling survival, resighting and movement probabilities.

Table S4. Goodness-of-fit tests of the general JMV model by sex and cohort and applied to the resighting data of greater flamingos born in the Camargue. *ĉ* denotes the overall overdispersion factor estimate. F=female; M=male.

Table S5. Modelling male resighting probability *p* with survival probability modelled as 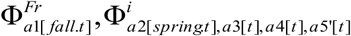 and movement probability modelled as 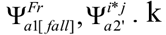, number of estimated parameters. Models are sorted by ΔAIC values.

Table S6. Modelling female resighting probability *p* with survival probability modelled as 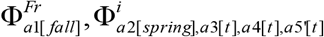 and movement probability modelled as 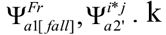, number of estimated parameters. Models are sorted by ΔAIC values.

Table S7. Annual survival probabilities (± SE) for greater flamingo chicks from semestrial survival probabilities estimated from model M3 and F1 for males and females respectively.

